# Quantifying antimicrobial access and practices for paediatric diarrheal disease in an urban community setting in Southeast Asia

**DOI:** 10.1101/228635

**Authors:** Le Thi Quynh Nhi, Ruklanthi de Alwis, Phung Khanh Lam, Nguyen Nhon Hoa, Nguyen Minh Nhan, Le Thi Tu Oanh, Dang Thanh Nam, Bui Nguyen Ngoc Han, Hoang Thi Huyen, Vu Thuy Duong, Lu Lan Vi, Bui Thi Thuy Tien, Hoang Thi Diem Tuyet, Le Hoang Nha, Guy Thwaites, Do Van Dung, Stephen Baker

## Abstract

Antimicrobial-resistant infections are increasing across Asia. Aiming to evaluate antimicrobial access and practices in Ho Chi Minh City (HCMC) of Vietnam, we mapped pharmacy locations and used a simulated client method to calculate antimicrobial sales for paediatric diarrheal disease. We additionally evaluated healthcare choices for parents and caregivers when their children experienced diarrhoea. District 8 (population 396,175) of HCMC had 301 pharmacies (one for every 1,316 people), with a density of 15.8 pharmacies/km^2^. A wide range of different treatments (n=57) were sold for paediatric diarrheal disease, with 8% (3/37) and 22% (8/37) of the sampled pharmacies selling antimicrobials for watery and mucoid diarrhoea, respectively. Despite the apparent abundance of pharmacies, the majority of caregivers chose to take their child to a specialized hospital, with 81% (319/396) and 88% (347/396) of responders selecting this as their first, second, or third choice for watery and mucoid diarrhoea, respectively. Lastly, by combining denominators derived from caregiver interviews and diarrheal incidence figures, we calculated that 16% (2,359/14,427) of watery or mucoid diarrhoea episodes of the District 8 population aged 1 to <5 years would receive an antimicrobial for diarrhoea annually, but antimicrobial prescribing was almost ten times greater in hospitals than in the community. Our novel mixed-methods approach found that, whilst antimicrobials are commonly available for paediatric diarrhoea in the community of HCMC, usage is greater in hospitals. The observed non-standardized approach to diarrheal treatments is indicative of poor recommendations. We advocate better guidelines, training and dissemination of information regarding antimicrobials and their use in this location.

## Introduction

Antimicrobial resistance (AMR) is rapidly becoming a public health issue of global concern.^1^ In the past decade, many of the key antimicrobials groups that we have come to rely in human medicine are in the process of losing, or have already lost, their effectiveness in treating infections caused by important bacterial pathogens ^2,3^. AMR is a complex scientific, political, economic, and social issue, and whilst developed countries are leading research efforts to tackle the problems, the greatest impact of AMR is currently being felt in low-middle income countries (LMICs).^4^–^6^ The AMR issue is magnified in LMICs because the principal driver of AMR is antimicrobial usage. Unrestricted access to low-cost, antimicrobials combined with a high burden of infectious disease, limited capacity for diagnostics, and large urbanizing populations demanding better medical support means that AMR bacteria are prospering across the LMICs in Asia.^7,8^

One of the main issues influencing AMR is the indiscriminate usage of antimicrobials for treating common human diseases, of which diarrheal disease is likely a major contributor. It was estimated that the worldwide burden of diarrhoea was >1.7 billion episodes in 2010, with the vast majority of these episodes arising in LMICs.^9^ The etiological agents of diarrhoea are broad (viruses, bacteria, and parasites) and can induce an array of symptoms, the more severe of which may necessitate an antimicrobial. Currently, the WHO recommends oral rehydration solution (ORS), zinc, and an antimicrobial for those with more severe diarrhea.^10,11^ However, a large disease burden, combined with a lack of financial and diagnostic resources means that the causative agent in LMICs is rarely or never identified.^12^ This lack of a confirmative diagnosis results in the empirical use of antimicrobials in the community or a healthcare facility. However, many of those receiving antimicrobials for diarrhoea may not require them ^13^, as the disease is generally self-limiting, is most likely to be of viral aetiology,^14^ and in LMICs in Asia many diarrheagenic bacteria are resistant to empirical antimicrobials.

Vietnam is a rapidly developing LMIC in Southeast Asia with a current population of >90 million people, which is estimated to be >100 million by 2020 ^15^. Ho Chi Minh City (HCMC) is the major financial centre in the south of the country and a typical urban, modernizing Asia megacity. An increasing population (>8 million people), a high infectious disease burden, and easy antimicrobial access represents an unparalleled opportunity for investigating AMR and antimicrobial usage. Diarrheal disease is common in HCMC, with the two main paediatric hospitals jointly admitting >20,000 children with diarrhoea per annum.^16^ Here, we aimed to assess access and practices related to antimicrobial sales by mapping pharmacy locations and using a simulated client method in an urban district of HCMC. Further, hypothesizing that antimicrobial usage for diarrheal disease is excessive in this setting, we measured the primary choices of healthcare services for the parents or caregivers of children with diarrheal disease in the community. By combining denominators derived from parents and caregivers interviews we were able to estimate the annual antimicrobial use for watery and mucoid diarrhoea in children aged 1 to <5 years living in central HCMC.

## Methods

### Ethics

Ethics approvals for the mapping of the pharmacies, pharmacy practices, and community surveys were provided through a minimal risk application by the Oxford University Tropical Research Ethics Committee (OxTREC approval 5110-16) and the local Institutional Review Board (IRB) of the University of Medicine and Pharmacy at HCMC (No. 220/DHYD-HD). Written informed consent was obtained from all participants over 18 years of age, or from parents and guardians if participant was under 18 years.

### Study setting and design

With an estimated population of >8 million people, HCMC is the largest city in southern Vietnam ^17^. The city is divided into five rural districts and 19 urban districts spread over an area of 19 km^2^; the urban districts had a mean population density of 13,394 people/km^2^ in 2016 ^17^. We selected District 8 in which to quantify access to vendors selling antimicrobials within HCMC. District 8 is located in the central part of the city and is divided into 16 wards (Figure S1), and had a population density of 22,522 people/km^2^ as of 2016 ^17^ (Table 1).

**Table 1.**
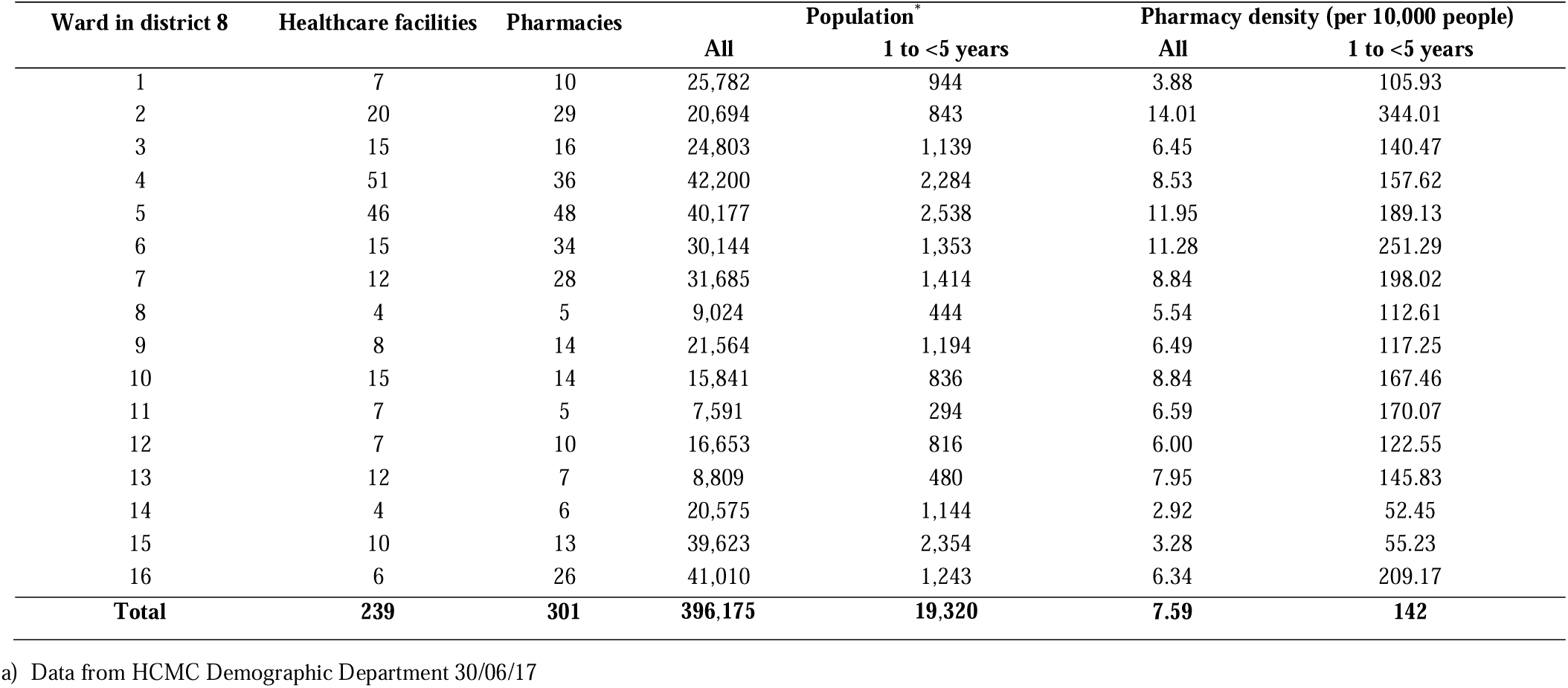
Pharmacies and healthcare clinics in the various wards of district 8 in Ho Chi Minh City

| Ward in district 8 | Healthcare facilities | Pharmacies | Population<sup>a</sup> |  | Pharmacy density (per 10,000 people) |  |
| --- | --- | --- | --- | --- | --- | --- |
|  |  |  | All | 1 to <5 years | All | 1 to <5 years |
| 1 | 7 | 10 | 25,782 | 944 | 3.88 | 105.93 |
| 2 | 20 | 29 | 20,694 | 843 | 14.01 | 344.01 |
| 3 | 15 | 16 | 24,803 | 1,139 | 6.45 | 140.47 |
| 4 | 51 | 36 | 42,200 | 2,284 | 8.53 | 157.62 |
| 5 | 46 | 48 | 40,177 | 2,538 | 11.95 | 189.13 |
| 6 | 15 | 34 | 30,144 | 1,353 | 11.28 | 251.29 |
| 7 | 12 | 28 | 31,685 | 1,414 | 8.84 | 198.02 |
| 8 | 4 | 5 | 9,024 | 444 | 5.54 | 112.61 |
| 9 | 8 | 14 | 21,564 | 1,194 | 6.49 | 117.25 |
| 10 | 15 | 14 | 15,841 | 836 | 8.84 | 167.46 |
| 11 | 7 | 5 | 7,591 | 294 | 6.59 | 170.07 |
| 12 | 7 | 10 | 16,653 | 816 | 6.00 | 122.55 |
| 13 | 12 | 7 | 8,809 | 480 | 7.95 | 145.83 |
| 14 | 4 | 6 | 20,575 | 1,144 | 2.92 | 52.45 |
| 15 | 10 | 13 | 39,623 | 2,354 | 3.28 | 55.23 |
| 16 | 6 | 26 | 41,010 | 1,243 | 6.34 | 209.17 |
| **Total** | **239** | **301** | **396,175** | **19,320** | **7.59** | **142** |

a) Data from HCMC Demographic Department 30/06/17

We conducted a cross-sectional study including three components, i) geographical mapping and of all pharmacies and healthcare facilities in District 8 of HCMC, ii) a pharmacy practice study to assess medication suggested by retail pharmacies for paediatric diarrhoea, and iii) a self-reported caregiver community survey on the management and antimicrobial treatment of paediatric diarrhoea.

### Geographical mapping of pharmacies in District 8

Addresses of all registered health clinics and pharmacies in District 8 were obtained from the HCMC Department of Health. Global Positioning System (GPS) coordinates of all pharmacies in District 8 were measured using hand-held electronic devices, such as mobile phones and tablets, and recorded using the geo-location tool EpiCollect5^18^.

### Pharmacy practice study

We conducted a pharmacy behaviour study using an unbiased observational technique for assessing dispensed medication called simulated client method or mystery shopper methodology ^19^–^22^. Two “mystery shoppers” with no official health training role-played as caregivers and approached pharmacies in Ward 5 of District 8 to seek medical advice and treatment for two scenarios of diarrhoea. *Scenario 1*: A mother attending the pharmacies in the study area to buy medication for a child that is 2 years of age. The child has had 4-5 loose stool episodes within the previous 24 hours. The child has no fever, and no blood or mucus in stools. *Scenario 2*: A mother attending the pharmacies in the study area to buy medication for a child that is 2 years of age. The child has had 4-5 loose stool episodes within the previous 24 hours. The child had a mild fever and mucus in the stool. No additional symptoms were provided if asked by the drug seller (i.e. no vomiting etc.) in both scenarios. Pharmacy-prescribed drugs were labelled anonymously and then were classified by a clinician and confirmed by a qualified pharmacist.

### Self-reported behaviour survey in the community

We performed a cross-sectional study in District 8 to capture the self-reported behaviour of parents and caregivers in the community in managing paediatric diarrhoea. The study population consisted of parents and caregivers of children (aged <5 years) residing in District 8, HCMC. We combined proportional sampling techniques including stratified random and simple random sampling. Firstly, a stratified random sampling technique was used to capture a representative sample of the identified population stratified by 16 wards and by age of the children. Secondly, a simple random sampling technique was applied to the currently registered children aged <5 years in the area to identify available individuals for the interviews. Local health authorities accompanied and assisted in community visits. A second home visit was made if caregiver was absent during the first visit. Those who had relocated from District 8 were replaced using a child of similar age chosen by random sampling. Mothers or caregivers were asked a short questionnaire regarding their choice of care if their child had diarrhoea in simulated scenarios similar to those described in the pharmacy behaviour study, this is provided in supplementary information. We surveyed 396 parents or caregivers.

### Geospatial analysis

Vietnam administrative boundaries were downloaded from Global Administrative Divisions Map (GADM). Vietnam population data rasters for 2015 (i.e. population per 0.01 Km^2^) were downloaded from WorldPop (100 m resolution), and satellite imagery of Vietnam was downloaded from ESRI basemaps. All geospatial mapping was conducted using ArcGIS v10.2 (Redlands CA, USA). Euclidean distance to the nearest pharmacy and kernel density plots for pharmacy density (i.e. pharmacies per 0.01 Km^2^) was estimated for District 8 using the “*Euclidean distance*” and “*Kernel*” tools in ArcGIS, respectively. Raster map depicting pharmacies per 10,000 people was computed by dividing the pharmacy density raster by the 2015 population raster using “*Raster Calculator*” in ArcGIS.

### Quantifying antimicrobial use for diarrheal treatment in the community and hospital

Here, antimicrobial usage for diarrheal disease is dependent on, a) the number of diarrheal episodes seeking medical treatment, and b) how likely antimicrobials are to be prescribed for diarrheal treatment (Figure 1). We calculated the annual number of diarrheal episodes amongst children aged 1-5 years in district 8 presenting at pharmacies as the product of, i) the total number of children aged 1-5 years in district 8, ii) the incidence rate of diarrhoea episodes per child-year, and iii) the proportion of parents or caregiver selected to firstly take their child to a pharmacy. To calculate the annual number of diarrheal episodes amongst children aged 1-5 years in district 8 presenting at hospitals, we multiplied i) the total number of children aged 1-5 years lived in district 8 and, iv) the incidence rate of diarrhoea episode presenting at hospital per child-year. The population i) was derived from recent census data from the HCMC Demographic department (30/06/17). The incidence rates ii) and iv) were estimated from a prospective diarrheal cohort conducted between 2014 and 2016 that actively followed 748 children in residing District 8 ^23^. The proportion of parents or caregivers seeking pharmacies as their first choice iii) was derived from the self-reported behaviour survey of parents and caregivers. To quantify the likelihood of prescribing antimicrobials for diarrheal treatment at a pharmacy or a hospital, we used data from the pharmacy behaviour study and the prospective cohort study ^23^, respectively. We estimated antimicrobial use for diarrheal treatment for both watery and mucoid diarrhoea. We generated each estimate with a corresponding 95% confidence interval (95CI), which takes into account uncertainty in the estimation of diarrhoea incidence only. The underlying figures described above are provided in Table S1.

**Figure 1.**
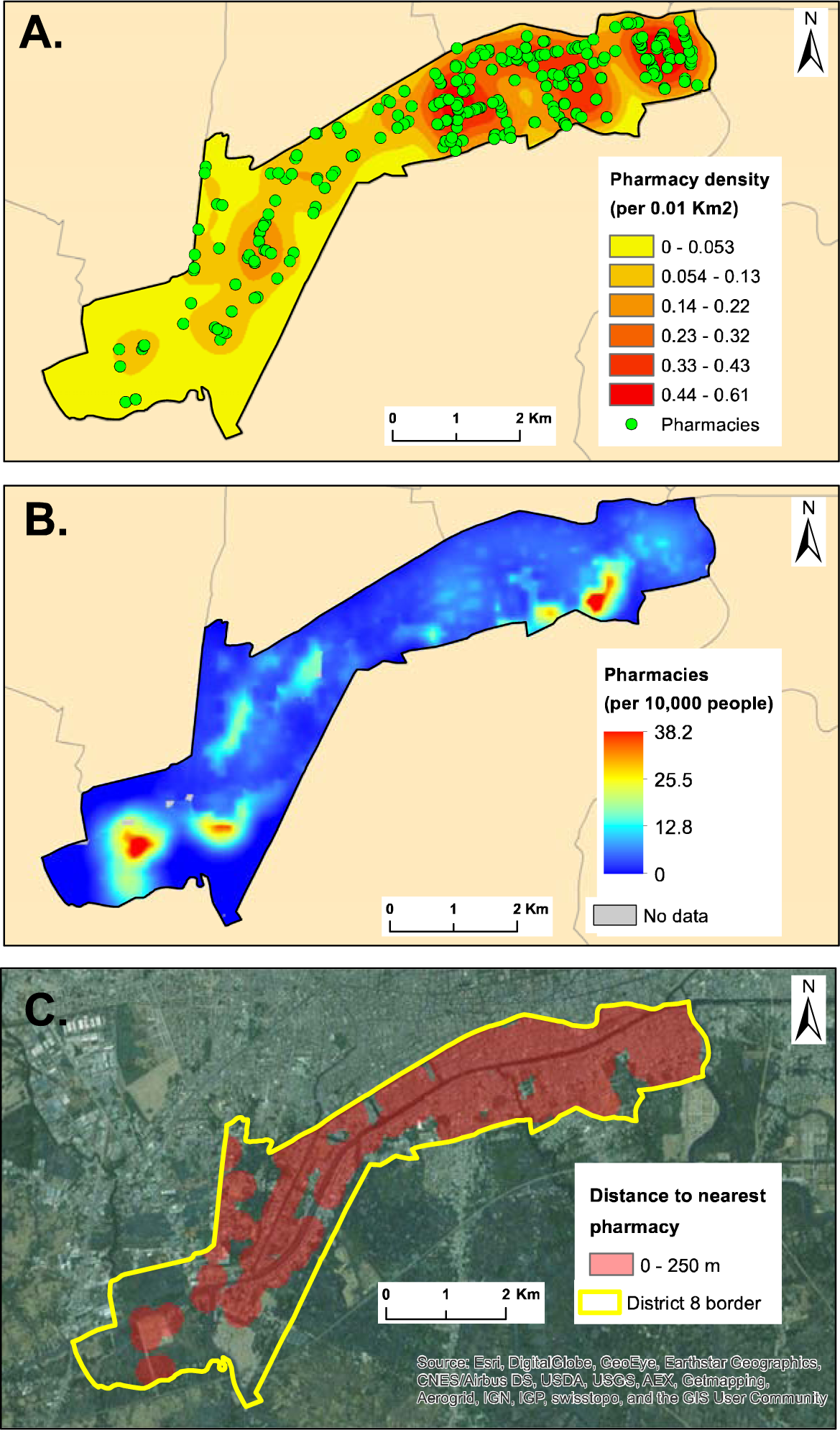
The geographical distribution of pharmacies in district 8 of Ho Chi Minh City Maps of district 8 in Ho Chi Minh City showing; A) the point locations of pharmacies and the kernel distribution of pharmacy shop density. Green points identify the pharmacy shops and pharmacy density is determined by colour intensity (see key). B) The distribution of pharmacy shops density per 10,000 inhabitants. The greater the intensity of red the higher the numbers of pharmacy shops per 10,000 people (see key, areas in grey are those with no data due to missing data in the population raster). C) The geographical areas in district 8 that are within 250m of a pharmacy. Red shaded areas are locations in with residents live with 250m of a location selling antimicrobials.

## Results

### The density and distribution of outlets selling antimicrobials in District 8 in HCMC

In September 2016 we mapped a total of 301 pharmacies and 243 private or public healthcare clinics in District 8 (Table 1 and Figure 1A). The distribution of pharmacies was not uniform and ranged from 0 to 61 pharmacies/km^2^, with a mean density of 15.7 pharmacies/km^2^. The majority of outlets were located on main thoroughfares in the northern portion of the district, specifically in wards 4, 5, and 6 (Figure 1A). The overall number of pharmacy outlets equated with 7.59 pharmacies/10,000 people, which corresponded with a pharmacy for every 1,316 people. The number of pharmacies within the population varied from 0 to 38.2 pharmacies/10,000 people, with four high density (>30 outlets /10,000 people) areas; two in the north and two in the south of the district (Figure 1B). The high density of pharmacy shops exemplified routine access to antimicrobials in this community, and equated with >75% of this population living within 250m of an outlet selling antimicrobials (Figure 1C).

### Pharmacy prescribing practices for diarrhoea disease in the community

We developed a simulated client approach and employed two “mystery shoppers” to visit the 48 pharmacy shops (37 were open at the time of the visits) in ward 5 of district 8 (Figure S1) and defined two different scenarios for paediatric diarrhoea. In the first scenario (watery diarrhoea), the number of prescribed medications ranged from one to five, with the majority (65%; 24/37) of pharmacies prescribing two different medications (Figure 2B). The median total cost for each pharmacy visit was 17,000VND (1USD= ~23,000VND); range 7,000 to 54,000VND. A total of 46 different types and brands of medications were sold, which could be broadly categorized into 11 classes; anti-motility, zinc, propulsive, anti-secretory, other, oral rehydration therapy (ORT), antimicrobial, antipyretic, unlabeled, adsorbent, and probiotic. Unlabeled treatments (sold without additional advice) were prescribed in 16% (6/37) of pharmacy visits, with probiotics (81%; 30/37) and adsorbents (68%; 25/37) being the most commonly sold treatments. Antimicrobials were prescribed by 3/37 (8%) of pharmacies (Figure 2A).

**Figure 2.**
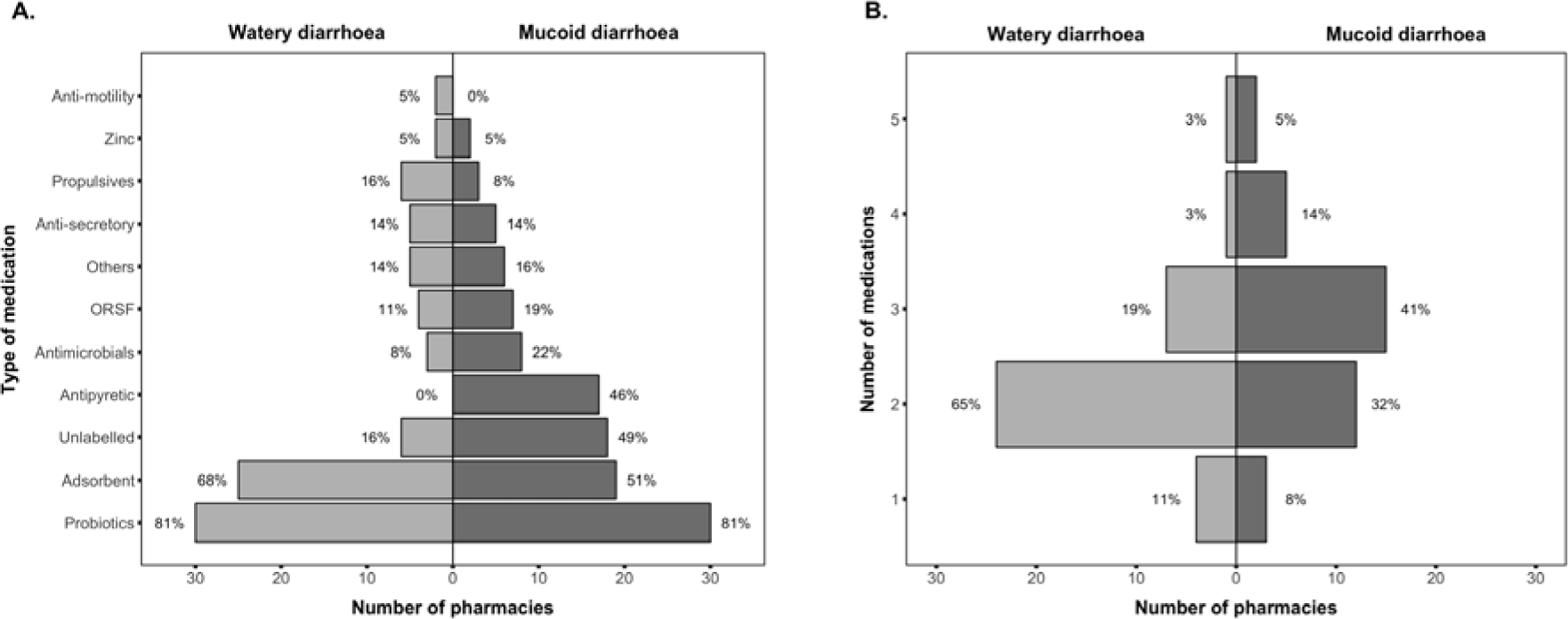
Medications sold by pharmacies in Ward 5 in District 8 for paediatric diarrhoea A) Rotated histogram showing the number and proportion (x-axis) of the 37 pharmacies visited during the “mystery shopper” simulated clients method selling different mediations (y-axis) for watery and mucoid diarrhoea, respectively. B) Rotated histogram showing the number and proportion (x-axis) of the 37 pharmacies visited during the “mystery shopper” simulated clients method selling between 1 and 5 medications (y-axis) for watery and mucoid diarrhoea, respectively.

In the second scenario (mucoid diarrhoea with fever), 53 different medications were sold by the 37 pharmacies; undefined/unlabeled medications were sold in 49% (18/37) of pharmacy visits (Figure 2A). Probiotics (81%; 30/37) and adsorbents (51%; 19/37) remained the most common medications prescribed, but the proportion of pharmacies prescribing an antimicrobial increased to 22% (8/37). There was also an increase in the number of mediations sold for mucoid diarrhoea in comparison to watery diarrhoea (Figure 2B), with a higher proportion of pharmacies selling a combination of three (41%; 15/37), four (14%; 5/37), or five (5%; 2/37) medications. Despite there being a greater number of medications sold, the cost to caregivers in this scenario was not substantially greater than with watery diarrhoea (median 18,000 VND; range 3,000-31,000 VND).

There was considerable disparity in the type of antimicrobials sold in the 37 sampled pharmacies, with pharmacies prescribing antimicrobials from five different classes: trimethoprim-sulphate, sulphonamides, fluoroquinolones, macrolides, and cephalosporins (1^st^, 2^nd^, and 3^rd^ generation). In a third scenario, when the mystery shoppers returned to the same pharmacy shops a week later and asked to buy ciprofloxacin, 100% (37/37) of the vendors sold the requested antimicrobial. This medication was reliably sold without dosing advice and without the caregiver having to provide information as to why the antimicrobial was required. Generic ciprofloxacin was sold in a range of different packaging from 14 different manufacturers in two different concentrations.

### Knowledge, attitudes, and practices regarding antimicrobial usage in caregivers

We next aimed to evaluate how the community in district 8 access antimicrobials and understand their knowledge, attitudes, and practices regarding antimicrobial usage. We interviewed 396 individuals that were either caregivers or parents of children aged <5 years residing in district 8. The demographic characteristics of these participants are summarized in Table S2. The majority (82%; 325/396) of those interviewed were woman and the median age was 39 years (range 19 to 87 years). A small fraction of those interviewed (10%; 39/381) indicated they had some background in healthcare. The majority (69%; 243/351) of interviewees described themselves as living in a household of average income (≥1,300,000 ≤9,000,000VND/household/month), with 21% (74/351) reported living in a poor household (<1,300,000VND /household/month). Most caregivers/parents (60%; 232/386) had ≥2 children living in the household. Many (59%; 231/394) reported that their children had received medication for a medical condition within the last month, of which approximately half (47%; 109/231) were reported to be an antimicrobial.

We found that most of parents and caregivers has limited knowledge about antimicrobials. Many assumed that antimicrobials could be generically used to treat coughing (46%; 181/396), fever (35%;137/396), colds (30%; 118/396), headaches (21%; 81/396), and diarrhoea (19%; 76/396). Approximately half (51%; 203/396) of the parents and caregivers agreed with the statement “some infections can be difficult to treat if there is antimicrobial resistance” and a 59% (233/396) agreed with the statement “resistance can occur if they do not take a sufficient dose”. In self-reported prescribing, more half of parents and caregivers (54%; 213/394) said they regularly purchased antimicrobials from pharmacies, with many of the (85%; 182/213) buying antimicrobials within the 30 days prior to the interview (Table S2).

### Diarrheal treatment seeking behaviour by caregivers in the community

Aiming to quantify antimicrobial usage for diarrheal disease in children resident in District 8, we additionally asked the parents and caregivers to highlight their top three preferences for where they would seek healthcare advice if their children had either of the two previously described diarrheal disease scenarios (watery or mucoid). We provided the responders eight different alternatives, which included pharmacies, private clinics, and specialized hospitals (Figure 3). In both scenarios the most common first choice was to treat their children at home. However, the most common cumulative option was taking their child to a specialized hospital, with 81% (319/396) and 88% (347/396) of responders selecting this as a first, second, or third choice for watery and mucoid diarrhoea, respectively. Visiting a local pharmacy as a first choice ranked third for watery diarrhoea and second for mucoid diarrhoea. Overall, visiting a pharmacy was a first, second, or third choice for 45% (180/396) and 34% (134/396) for caregivers with children with watery and mucoid diarrhoea, respectively (Figure 3). We found that that parents (rather than a caregivers) and those of younger age were more likely to take their children to a pharmacy than a hospital for watery diarrhoea. We also found that parents (rather than a caregivers) were significantly more likely to take their children to a pharmacy than a hospital for mucoid diarrhoea (Table 2)

**Figure 3.**
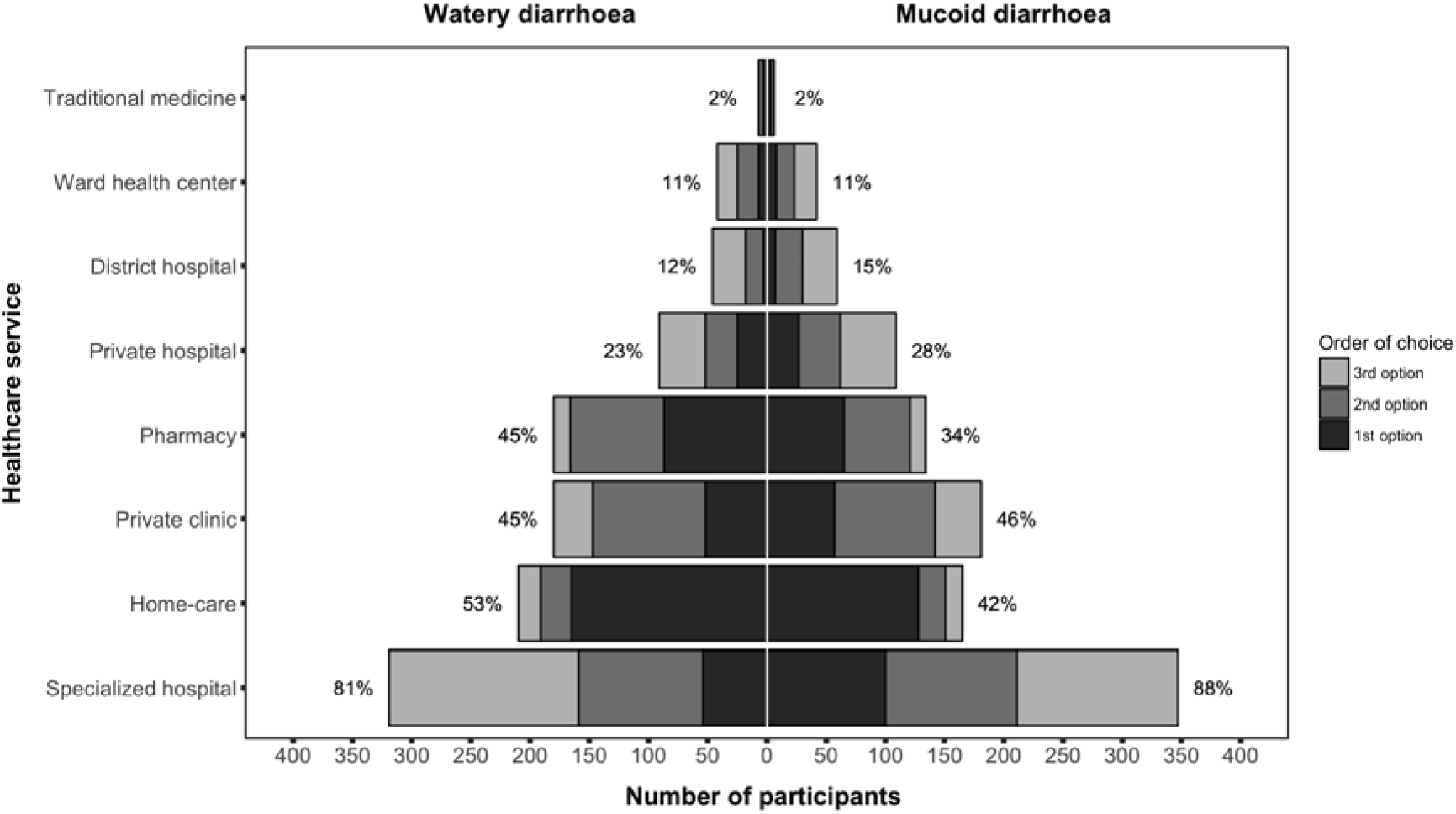
Healthcare utilization for paediatric diarrhoea by caregivers in District 8 Pyramid plot showing the number and proportions of 396 individuals caregivers or parents of children aged <5 years residing in District 8 (x-axis) selecting the various healthcare services (y-axis) as first, second, or third choices (shaded, see key) for watery and mucoid diarrhoea.

**Table 2.** The demographic characteristics of parents and caregivers who chose a pharmacy or a hospital as first choice for a child with diarrhoea

|  | Watery diarrhoea |  |  |  |  | Mucoïd diarrhoea |  |  |  |  |
| --- | --- | --- | --- | --- | --- | --- | --- | --- | --- | --- |
|  | n | Pharmacy first choice<br>(n=87) | n | Hospital first choice<br>(n=89) | p value <sup>a</sup> | n | Pharmacy first choice<br>(n=65) | n | Hospital first choice<br>(n=142) | p value <sup>a</sup> |
| Female sex | 87 | 73 (84%) | 89 | 72 (81%) | 0.693 | 65 | 55 (85%) | 142 | 113 (80%) | 0.448 |
| Role of parent (not caregivers) | 86 | 54 (63%) | 89 | 39 (44%) | **0.015** | 64 | 43 (67%) | 142 | 69 (49%) | **0.015** |
| Age in years (IQR) | 81 | 38 (30, 50) | 88 | 40 (32, 57) | **0.051** | 60 | 38 (31, 50) | 139 | 40 (32, 56) | 0.198 |
| Healthcare trained | 83 | 7 (8%) | 89 | 11 (12%) | 0.461 | 63 | 4 (6%) | 138 | 16 (12%) | 0.316 |
| Employment | 87 | 87/87 (100%) | 89 | 89/89 (100%) | 0.661 | 65 | 65/65 (100%) | 142 | 142/142 (100%) | 0.943 |
| Worker |  | 10 (11%) |  | 10 (11%) |  |  | 6 (9%) |  | 16 (11%) |  |
| Officer |  | 6 (7%) |  | 6 (7%) |  |  | 5 (8%) |  | 10 (7%) |  |
| Health staff |  | 2 (2%) |  | 6 (7%) |  |  | 1 (2%) |  | 6 (4%) |  |
| Business owner |  | 15 (17%) |  | 10 (11%) |  |  | 11 (17%) |  | 23 (16%) |  |
| Housewife |  | 35 (40%) |  | 34 (38%) |  |  | 29 (45%) |  | 56 (39%) |  |
| Other (retired) |  | 19 (22%) |  | 23 (26%) |  |  | 13 (20%) |  | 31 (22%) |  |
| Education ≤12 years | 87 | 75 (86%) | 88 | 68 (77%) | 0.171 | 65 | 55 (85%) | 141 | 115 (82%) | 0.695 |
| Monthly income <sup>b</sup> | 78 |  | 80 |  | 0.406 | 57 |  | 127 |  | 0.739 |
| ≤1.3M VND |  | 15 (19%) |  | 14 (18%) |  |  | 11 (19%) |  | 20 (16%) |  |
| 1.3-9M VND |  | 59 (76%) |  | 57 (71%) |  |  | 42 (74%) |  | 95 (75%) |  |
| >9M VND |  | 4 (5%) |  | 9 (11%) |  |  | 4 (7%) |  | 12 (9%) |  |
| Only 1 child | 84 | 36 (43%) | 84 | 37 (44%) | 1.000 | 65 | 21 (32%) | 134 | 61 (46%) | 0.091 |
| Only 1 child aged <5 years | 85 | 68 (80%) | 86 | 71 (83%) | 0.699 | 65 | 49 (75%) | 134 | 105 (78%) | 0.718 |
| Age of first child (years) | 87 | 3 (2, 4) | 87 | 2 (2, 4) | 0.486 | 65 | 3 (2, 4) | 140 | 3 (2, 4) | 0.553 |
| Infections in previous month | 86 | 57 (66%) | 86 | 44 (51%) | 0.063 | 65 | 43 (66%) | 139 | 73 (53%) | 0.071 |
| Medication in previous month | 87 | 60 (69%) | 87 | 51 (59%) | 0.207 | 65 | 45 (69%) | 141 | 83 (59%) | 0.167 |
| Antimicrobials in previous | 60 | 24 (40%) | 70 | 26 (37%) | 0.857 | 41 | 18 (44%) | 118 | 46 (39%) | 0.585 |

a) p values calculated using Fisher's exact test for categorical variables and the Mann-Whitney U-test for continuous variables. Significant variable are highlighted in bold
b) Monthly income <1,300,000VND/month is determined to be poor and below average (as suggest by the Vietnamese government) <sup>46</sup>; we considered >=9,000,000VND/month a good monthly income and above average

**Table 3.** The burden of diarrhoea and antimicrobial treatment for diarrhoea in children aged 1 to 5 years in District 8

| Diarrhoea type | Diarrhoea episodes years per year (95% CI) |  |  | Episodes with antimicrobial treatment per year (95% CI) |  |
| --- | --- | --- | --- | --- | --- |
|  | Total | Attending pharmacy a | Attending hospital | Pharmacy | Hospital |
| Watery | 10,355 (8,945-11,765) | 2,275 (1,965-2,584) | 2,163 (1,622-2,724) | 184 (159-209) | 898 (673-1,130) |
| Mucoid | 4,057 (2,975-5,158) | 667 (489-848) | 2,241 (1,603-2,878) | 144 (105-183) | 1,462 (1,046-1,878) |
| Total | 14,427 (12,636-16,219) | 2,945 (2,585-3,305) | 4,404 (3,560-5,249) | 329 (282-375) | 2,359 (1,884-2,834) |

a) The 95% CIs did not take into account uncertainty in estimating % of attending pharmacy and % of receiving antimicrobials.

### Quantifying antimicrobial use for diarrheal treatment

We lastly aimed to calculate antimicrobial usage in the community and in hospitals for diarrheal disease in children aged between 1 and 5 years in district 8 of HCMC. Using data from a diarrheal disease cohort in district 8 we estimated the annual incidence of watery and mucoid diarrhoea in children aged 1 to 5 years to be 536 (463-609) and 210 (154-267)/1,000 child years of observation, respectively. We extrapolated these data to estimate the total number of diarrheal disease episodes in district 8 in this age group annually, which equated with 10,355 (8,945-11,765) and 4,057 (2,975-5,158) episodes of watery and mucoid diarrheal per year, respectively (Table 4). We found from the diarrheal treatment seeking behaviour interviews that paediatric watery diarrhoea were more likely to present at a pharmacy than a hospital, whereas mucoid diarrhoea cases was more likely to present at a hospital. Using data generated from a diarrheal cohort study and the simulated client method, we estimated the number (95% CI) of diarrhoea episodes in children aged 1 to 5 years that would receive antimicrobial treatment from a pharmacy in district 8 to be 184 (159-209) per year for watery diarrhoea and 144 (105-183) per year for mucoid diarrhoea. In those attending a hospital these numbers were estimated to be approximately five times higher (898 (673-1,130) per year) for watery diarrhoea and approximately ten times higher (1,462 (1,046-1,878) per year) for mucoid diarrhoea.

## Discussion

Antimicrobial resistant infections are increasingly throughout Asia, and are likely being driven in part by ease of access to antimicrobials. This factor is likely more relevant in a setting with a high burden of infectious diseases, low diagnostic capacity, and a large urbanizing population ^24^–^26^. Vietnam is the world’s fifteenth most populous country, with >96 million people in 2017 and urbanization increased from 28% in 2014 to 34.9% in 2017 ^27^. Antimicrobial dispensing in Vietnam is known to be poorly regulated, leading to inappropriate prescribing and self-medication for a number of common infections, including acute respiratory infections ^28,29^. It is also known that antimicrobials usage is also excessive for paediatric diarrheal disease, which is a common in Vietnam, and among the top motives for seeking healthcare facilities in HCMC ^12,30–32^. However, it was unknown whether antimicrobial usage for treating paediatric diarrheal disease was more prevalent in the community or a hospital setting. Here, we report simple access to community outlets selling antimicrobials in an urban district of HCMC. Our data suggest that antimicrobial usage for paediatric diarrheal diseases is excessive in this setting, but surprisingly, antimicrobial usage for paediatric diarrheal diseases it almost ten times more common in hospitals than in the community.

The general density of pharmacies in the community in HCMC is high (7.59/10,000 people) and outnumbers the number of pharmacy staff that have been trained at any level (assistant or university degree) in this city (5.98/10,000 people)^33^. This density of pharmacies in HCMC is reflective of the general pattern in LMICs, for example Sabde *et al.* reported a comparative distribution in an urban of India (5.84/10,000 people)^34^. Alarmingly, the number of pharmacy outlets in HCMC is more than double the mean density of pharmacies in other areas in the world, such as Europe (3.06/10,000 people) ^35^, Southeast Asia (3.02/10,000 people) ^35^, three times higher than that of some countries in Western Pacific (2.28/10,000 people) ^35^, and the USA (2.11/10,000 people)^36^, and six times higher than that in parts of Africa ^35^. Ease of access to retail pharmacies may improve equity in the use of medication, but allows access to drugs that should be better controlled, especially when pharmacy staff are under qualified and/or fail to follow appropriate guidelines.

Our study demonstrated the common vending of antimicrobials without prescription for paediatric diarrheal disease in the community in HCMC. However, this practice varied according to the reported severity of the disease, in which less severe symptoms were associated with fewer dispended antimicrobials. Notably, we estimated that antimicrobials sales for paediatric diarrheal disease in the community in HCMC was lower than that in Ethiopia (26%; 58/223) ^20^, and other Asian countries, such as Pakistan (14%) ^37^, and Thailand (52%) ^38^. The low proportion of antimicrobials dispensed for acute watery diarrhoea in HCMC (8%, 3/37) is complementary with a declining trend in antimicrobials use for watery diarrhoea in Vietnam. Available data suggests that antimicrobials dispensed for acute watery disease has decreased over the last 20 years, from 45% in 1997 ^39^, to 14% in 2013 ^40^, to 8% (3/37) in our study). This trend may result from a substantial improvement in practices in pharmacies to alleviative antimicrobial prescribing and the impact of resistance, although antimicrobials remain among the most inappropriate drugs dispensed from retail pharmacies in Vietnam ^41^. It is additionally worth noting that previous studies suggested that antimicrobial usage is much greater in rural areas than in urban areas ^29^. Therefore, our results from an urban setting may underestimate the extent of the issue from other locations in and around HCMC and across the country.

Our major finding was that antimicrobials usage for paediatric diarrheal in HCMC is more common is hospitals than in the community. We predicted that the diarrheal disease management, including antimicrobials dispensing, in the community is poor in HCMC, which is similar to a number of other areas in the world ^37–39,42,43^.We found that the total number of diarrheal episodes that would receive an antimicrobial in the hospital was five to ten times higher than that from retail pharmacies; this observation was contrary to our hypothesis. Our findings are consistent with a recent finding in this setting, where 85% of bloody and mucoid paediatric diarrheal patients admitted to hospital were prescribed antimicrobials ^44^. While one-third of antimicrobials prescribed in hospitals may be considered inappropriate ^45^, this practice could enhance associated with AMR. A potential explanation for our finding is that children with diarrheal disease attending a specified hospital are generally considered to be the more serious cases. Therefore, doctors are inclined to prescribe first-line therapies to prevent disease progression and to satisfy parent expectations for treating the infection with the best available medication. It is apparent that, in a setting with limited-to-no diagnostic testing, that this clinical approach is thought to be optimal for preventing complication and saving lives. We suggest that antimicrobials are used an as “insurance policy” rather that as a therapeutic treatment option. There is a large demand for hospital care in HCMC and doctors have limited time to make clinical decisions and this approach is may more effective and less time consuming. Therefore, there is a clear requirement for tests that can identify children at the more severe end of the disease spectrum and that can simply identify between a viral and a bacterial infection.

Our study has limitations. First, our data extrapolation may contain a degree of error as we compiled data from several different studies conducted in HCMC. Second, although we approached every pharmacy in one busy ward of the district, the number of pharmacies sampled was relatively small. Therefore, our findings may not represent the entire practice of antimicrobial dispensing in pharmacies in urban areas of HCMC. Third, aiming to assess antimicrobials dispensing only, we did not record or evaluate the retailer’s consulting abilities for using these drugs, which may reflect antimicrobial handling practice in the community. Lastly, our combined studies focused on diarrheal diseases in young children only, we recognize that our approach limited the understanding of antimicrobial usage in other populations and for other presentations in this location. Notwithstanding these limitations, to our knowledge this is the first study that has used a mixed-methods approach to quantify pharmacy density in urban Vietnam. Through this combined approach, we have described the profile of antimicrobials usage for diarrheal disease treatment in an urban area in an LMIC. Our work is important for the assessment of antimicrobial access in the community in Vietnam, and will inform practices to public health organizations at local and national levels that require a better understanding the role of pharmacy retailers in managing AMR.

We conclude that the pharmacy retailer density is extremely dense in urban HCMC, which is associated with straightforward antimicrobial access in the community. However, antimicrobial usage for diarrhoea disease in children is more common in hospitals than through community pharmacies. Given these findings, combined with a general low level of knowledge regarding antimicrobial and the self-reporting behaviour parents and caregivers, we predict a high burden of antimicrobial overuse in the community in this location. We propose further applied research to improve the knowledge, attitudes, and practice of parents and caregivers in terms of antimicrobial prescribing and AMR. We additionally advocate better education, training, and guidelines for antimicrobial usage and AMR for all health professionals with the authority to prescribe antimicrobials in Vietnam. Lastly, rapid diagnostic tests at the first point of care contact are essential for optimizing antimicrobial therapy in the community.

---

**Table S1.** Denominators for measuring antimicrobial usage for diarrhoea in community and hospital settings

| **Diarrhoea type** | **Incidence of diarrhoea episode per person-years (SE) <sup>a</sup>** | **Number of children 1-5 years old** | **% Attending Pharmacies** | **Incidence of diarrhoea episode attending hospital per person-years (SE)** | **% antimicrobials Pharmacies** | **% antimicrobials Hospitals** |
| --- | --- | --- | --- | --- | --- | --- |
| Watery | 0.536 (0.037) | 19,320 | 87/396 (22%) | 0.112 (0.015) | 3/37 (8%) | 66/159 (42%) |
| Mucoid | 0.210 (0.029) | 19,320 | 65/395 (16%) | 0.116 (0.017) | 8/37 (22%) | 107/164 (65%) |

a) SE: standard error

**Table S2.**
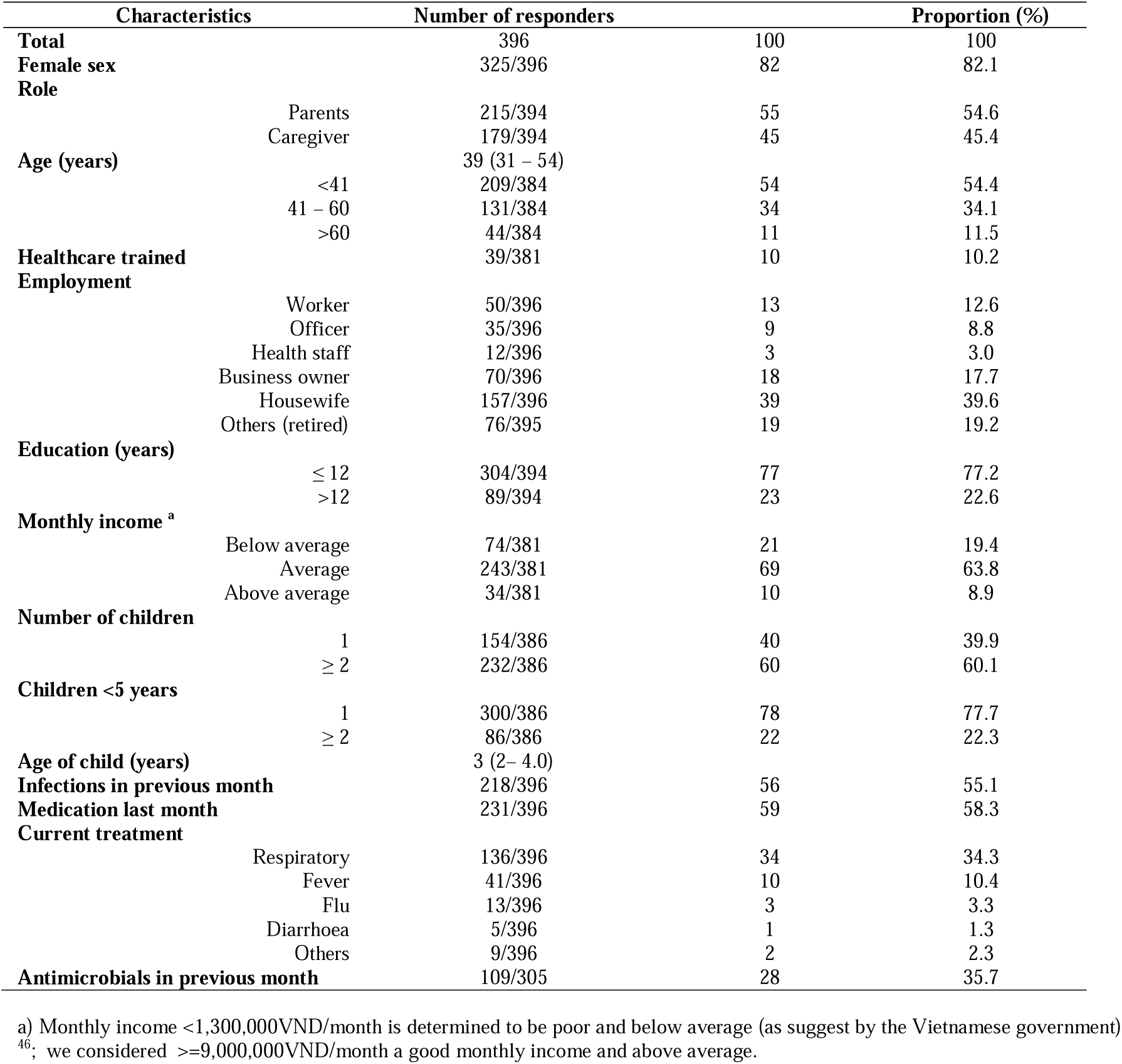
The demographic characteristics of parents and caregivers in the community behaviour survey

**Table S3.**
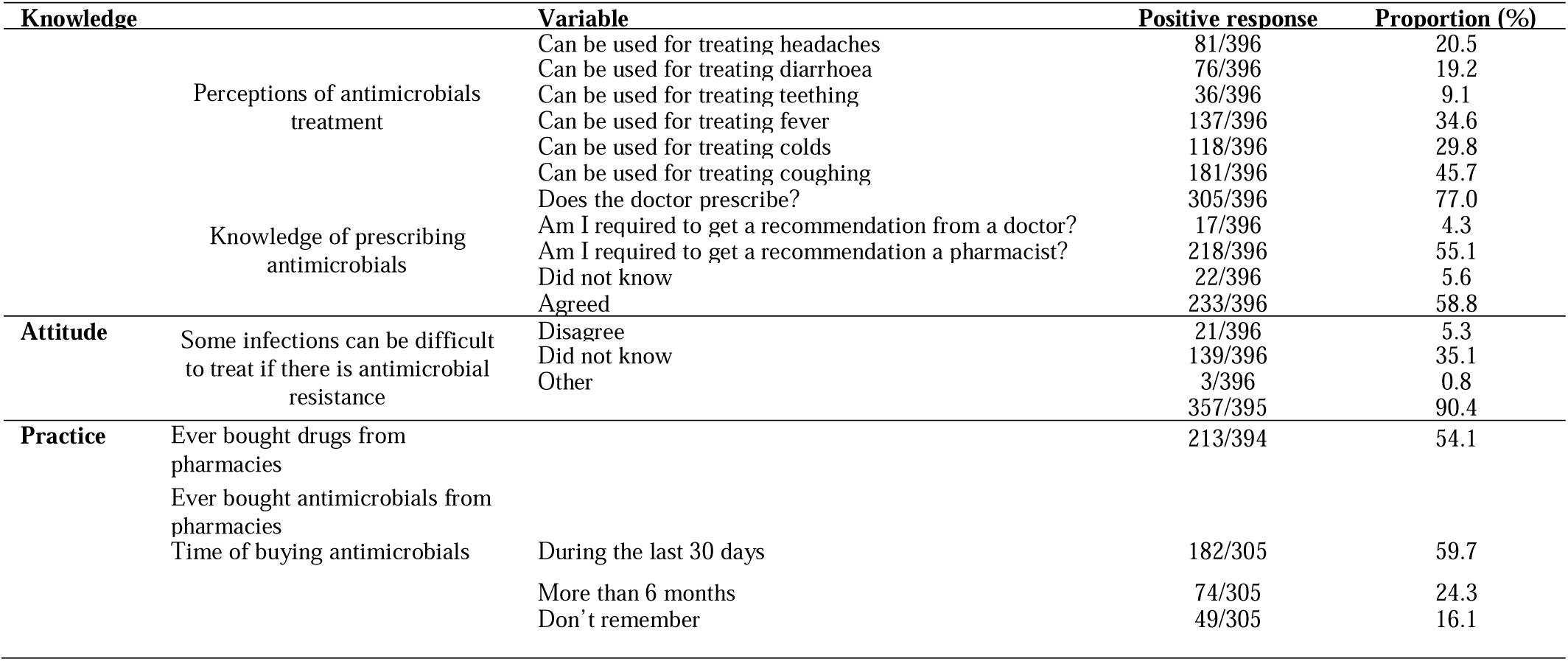
Knowledge, attitudes, and practice of parents and caregivers toward antimicrobials and antimicrobial resistance

**Figure S1.**
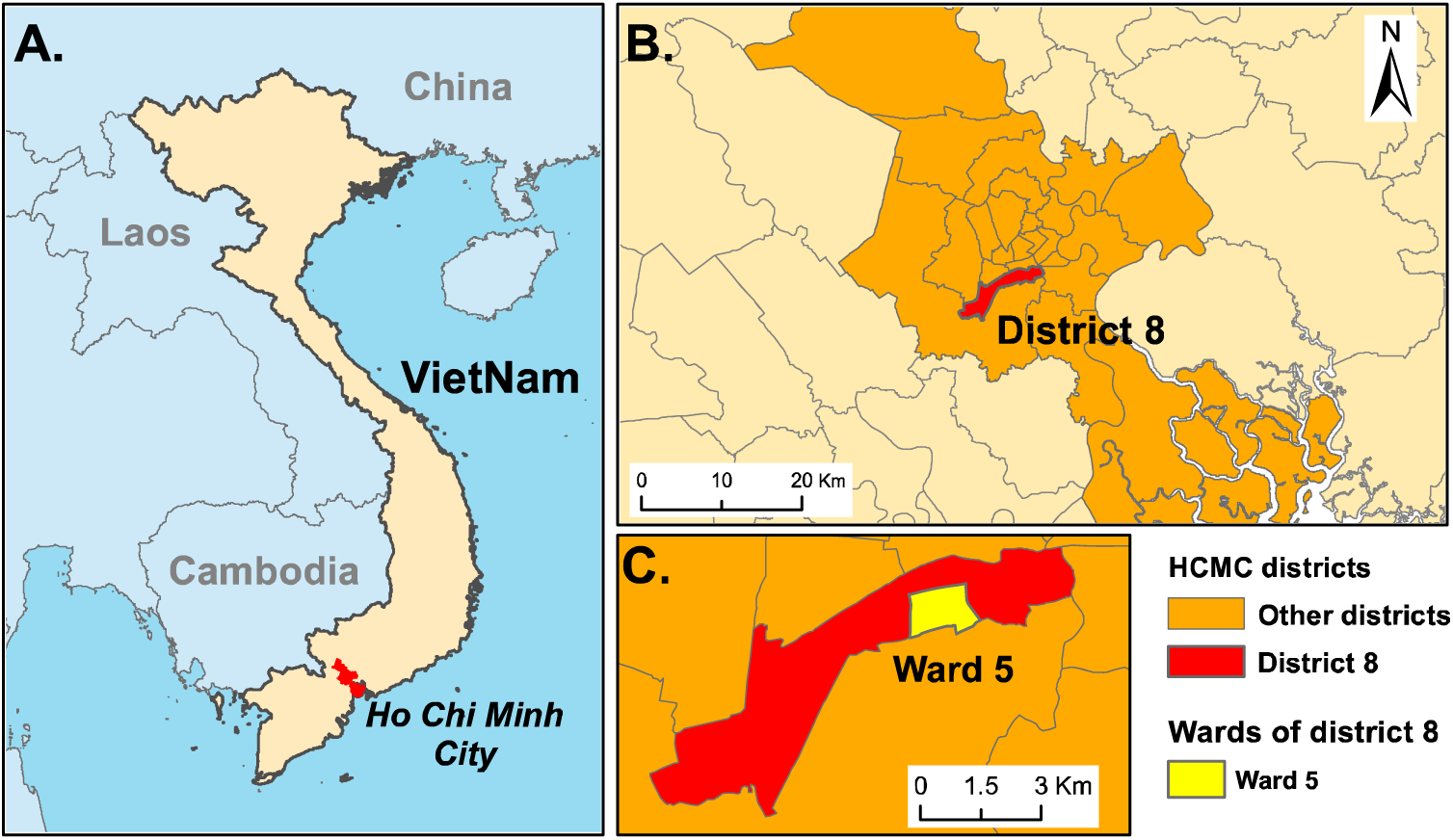
The selected geographical location for this study A) Map of Vietnam showing the location of the greater Ho Chi Minh City area. B) Enhanced map of Ho Chi Minh City showing the districts C) Enhanced map of district 8 of Ho Chi Minh City highlighting ward 5 as the location for the “mystery shopper” simulated clients method in yellow.

